# Integrative Multi-Scale Network Simulation for Precision Drug Repurposing with PETS

**DOI:** 10.1101/2025.02.24.639994

**Authors:** Kevin Song, Jake Y. Chen

**Author notes:** Corresponding author: Kevin Song.

## Abstract

Drug repurposing offers a promising strategy to accelerate therapeutic discovery for complex diseases; however, conventional approaches often overlook the multi-scale, dynamic nature of drug-induced perturbations. In this study, we introduce PETS (Predictive Evaluation of Therapeutic Simulations), an innovative *in silico* framework that synergistically integrates curated pathway models with multi-layer network propagation, adaptive dampening, and memory mechanisms to simulate drug–gene interactions with high fidelity. By propagating perturbations through tissue- and cell-specific protein–protein interaction networks via iterative state updates, PETS robustly captures both proximal and distal regulatory feedback, enabling the identification and ranking of drug candidates capable of reversing disease-specific gene expression signatures. Validation against the SigCom L1000 dataset demonstrates that PETS achieves lower mean squared errors and higher predictive consistency compared to existing methods across models of Alzheimer’s disease, AD-expressing neuroblastoma, and glioblastoma multiforme. Moreover, PETS concurrently quantifies on-target therapeutic efficacy and potential off-target toxicity, providing mechanistic insights that are critical for candidate prioritization. Looking ahead, the integration of patient-specific multi-omics data and dynamic network topologies is expected to further enhance the translational impact of PETS. Our framework represents a significant advancement in the computational prediction of drug–gene perturbation effects, positioning it as a cornerstone tool for precision medicine and preclinical drug discovery.

## 1 Introduction

Drug repurposing has emerged as a transformative paradigm in modern drug discovery, offering significant reductions in both development time and financial risk compared with traditional pipelines. Conventional drug development spans 12–15 years and can require investments on the order of 2–3 billion dollars [1; 2]. With nearly 90% of drug candidates failing during clinical trials due to safety or efficacy issues [4; 5], repurposing compounds with established safety and pharmacokinetic profiles presents a pragmatic strategy that not only accelerates time-to-market but also democratizes innovation for academic and smaller pharmaceutical entities [1; 3].

Advances in large-scale data generation and computational modeling have further expanded the scope of drug repurposing. Early efforts—ranging from mining electronic health records (EHRs) and clinical trial outcomes to applying guilt-by-association and phenotypic matching techniques (e.g., Connectivity Map)[8; 12; 13]—have been instrumental in identifying potential drug–disease associations. Moreover, the evolution of network medicine has reframed disease as perturbations within specific modules of the human interactome, where aligning known drug targets with disease-specific molecular signatures offers a mechanistic rationale for repurposing[6; 7]. Despite these advances, many existing computational methods fall short: they often rely on static representations of complex biological networks, lack detailed mechanistic insights, and do not seamlessly bridge the gap between computational predictions and clinical outcomes [16; 17; 18; 19].

A major step forward has been the integration of pathway models with drug–gene perturbation simulations. When drugs interact with their targets, they trigger cascades of molecular events that ultimately alter gene expression profiles [20; 21]. Recent simulation frameworks now emphasize identifying “influential genes” whose perturbation can drive broad network responses. These approaches utilize expansive databases—such as LINCS, DREAM Challenge, and CMap—to capture both immediate and delayed cellular responses using steady-state and time-series analyses [22; 23; 24; 25]. Complementary mathematical models—including Boolean networks, fuzzy logic systems, integer linear programming, and ordinary differential equations—offer quantitative insights into the temporal dynamics of signal transduction and regulatory network behavior [26; 27; 28; 29; 30; 31; 32]. Validation methods have similarly evolved, with modern protocols assessing model performance over a range of gene disruption levels and biological contexts [34; 35; 36; 37; 38].

Yet, challenges remain. The immense complexity of biological systems—characterized by tens of thousands of genes and an astronomical number of potential drug-like compounds [39; 40]—renders exhaustive experimental testing impractical. Moreover, many models simplify the intricate interplay and cross-talk among gene modules, often neglecting the temporal and spatial dynamics that are crucial for accurate predictions [42; 43; 44]. Clinically, while in silico predictions are promising, a significant translational gap persists; extensive preclinical testing and clinical validation are needed to bridge computational forecasts with patient outcomes [45]. Emerging strategies, such as integrating digital therapeutics with repurposed drugs, offer promising avenues to enhance treatment efficacy and monitoring [46; 47; 48].

In response to these limitations, we introduce the Predictive Evaluation of Therapeutic Simulations (PETS) framework—a novel, integrative approach that unites mechanistic pathway modeling with advanced network propagation algorithms. As depicted schematically in Fig.1, PETS constructs detailed, disease-specific pathway models that encapsulate curated molecular interactions and regulatory feedback loops. A key innovation within the PETS framework is its ability to simulate the dynamic evolution of cellular networks. Two novel components—an *adaptive dampening* mechanism and a *network memory* term—address the challenge of feedback regulation. The adaptive dampening mechanism modulates simulation intensity to prevent runaway oscillations, while the network memory term integrates historical perturbation data to capture delayed and cumulative cellular responses. Together, these features provide a more realistic depiction of drug-induced dynamics over time.

**Figure 1.**
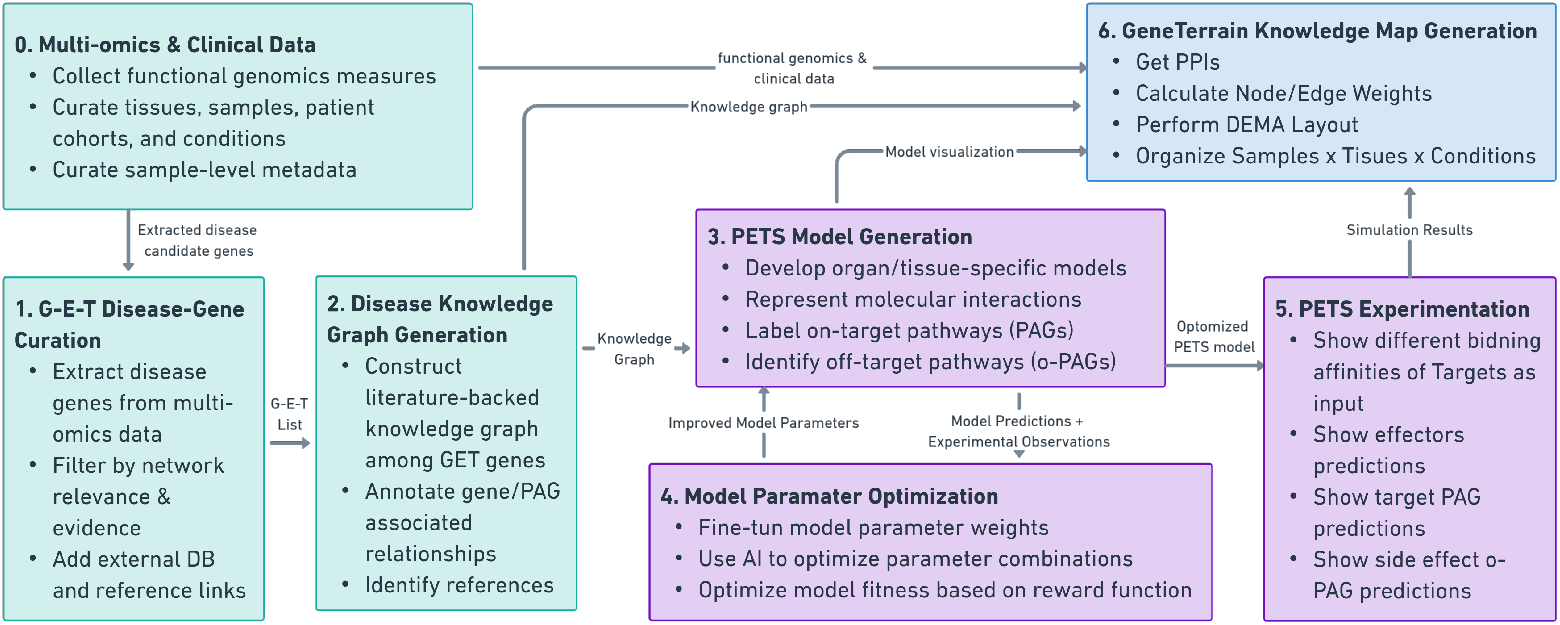
Overview of a programmable medicine framework that integrates multiomics and clinical data with literature-backed disease knowledge graphs, the PETS modeling pipeline, and GeneTerrain analysis. The process begins with diseasegene curation of gene sets encompassing genetic variants, differential expression, and drug targets (Steps 0–1) and knowledge graph construction (Step 2), followed by PETS model generation (Step 3) and parameter optimization (Step 4). The optimized models are then used for PETS experimentation (Step 5) to predict novel therapeutic targets, culminating in GeneTerrain knowledge map generation (Step 6) for comprehensive visualization of multiscale disease mechanisms and drug effects. This integrated approach aims to refine target discovery, guide drug repurposing, and accelerate clinical translation.

Programmable medicine is an emerging paradigm that utilizes advanced computational models, multi-omics data, and precise biological insights to tailor therapeutic strategies for individual patients or populations. The approach aims to enable dynamic, data-driven decision-making by incorporating diverse information sources—such as genomic, transcriptomic, proteomic, and clinical data—into sophisticated models of disease and treatment response. PETS (Predictive Evaluation of Therapeutic Simulations) is a critical component of this framework, offering an innovative solution for drug repurposing and target discovery. By integrating disease-specific pathway models, network propagation algorithms, and adaptive feedback mechanisms, PETS can predict therapeutic effects with enhanced precision and mechanistic insight. This capability makes it especially valuable in programmable medicine, where personalized treatments can be designed based on the specific molecular characteristics of a patient’s disease. Through its integration into a broader ecosystem of literature-backed knowledge graphs and tools like GeneTerrain analysis, PETS enables the creation of highly accurate, context-specific treatment strategies aimed at optimizing patient outcomes.

## 2 Materials and Methods

### 2.1 Multi-Scale Simulation Framework for Alzheimer’s Disease

To capture the complex, multi-scale pathophysiology of Alzheimer’s disease (AD), we developed an *in silico* simulation framework that integrates tissue- and cell-specific network dynamics. The framework models drug perturbations within an AD-relevant protein–protein interaction (PPI) network using an iterative state-update scheme. Implemented in Python, the algorithm combines multi-layer network propagation, standardized data preprocessing, adaptive dampening to mitigate oscillatory artifacts, and a composite loss function for robust parameter calibration.

### 2.2 Data Assembly and Preprocessing

A curated set of AD-relevant genes was obtained from PAGER (PAG ID: WIG000469) and augmented with regulatory interactions from the SIGNOR database. Let

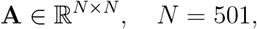

denote the PPI network, where each element *A*_*ij*_ represents both the confidence and sign of the regulatory interaction from protein *i* to protein *j*. For numerical stability, the network is normalized as

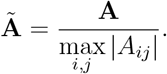

Ground-truth drug–gene perturbation data were sourced from the SigCom L1000 platform and encoded as a drug vector

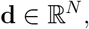

where nonzero entries denote direct drug–protein interactions. The drug vector is standardized (zero mean, unit variance) and used to initialize the state:

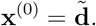

Additional data include AD-specific expression changes *E*_*i*_ (from Exp.csv) and protein importance weights *w*_*i*_ (from BEERE and WINNER, stored in Rp.csv). These data are incorporated into subsequent efficacy scoring and network modulation steps.

### 2.3 Network Enhancement via Multi-Layer Propagation

To capture higher-order interactions, multi-layer representations of the normalized adjacency matrix are computed as follows:

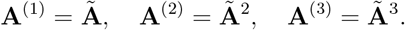

Each layer is scaled by confidence coefficients *c*_1_, *c*_2_, *c*_3_ and weighted by corresponding layer weights *α*_1_, *α*_2_, *α*_3_. Defining

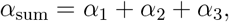

the enhanced network is constructed as

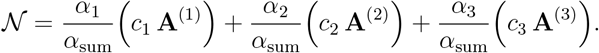

Protein importance weights *w*_*i*_ are incorporated by scaling with a factor *s*, where

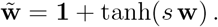

The scaled weights are applied element-wise to 𝒩, and the resulting entries are clamped to the interval [−1, 1].

### 2.4 Iterative State Update

The protein activation state vector **x**^(*k*)^ ∈ ℝ^*N*^ evolves over discrete time steps *k* = 1, …, *T* (with *T* = 300). At each iteration, the enhanced network 𝒩 is applied to the current state, and drug effects are emphasized through a *Boosting Vector*. In this framework, the boosting vector is defined for each protein *i* as

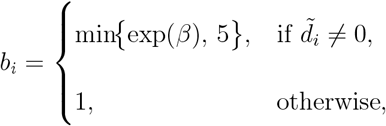

where *β* is a parameter controlling the amplification of nonzero drug perturbations. The state update is then given by

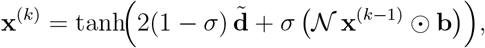

where *σ* ∈ [0, 1] balances the influence of the drug input versus network feedback, and ⊙ denotes element-wise multiplication.

As the simulation progresses, two additional mechanisms are incorporated. First, *adaptive dampening* is used to prevent oscillatory behavior. Inspired by negative feedback in biological systems, a dampening factor *δ*(*k*) is computed for *k* ≥ *k*_damp_ (with *k*_damp_ = 20) as

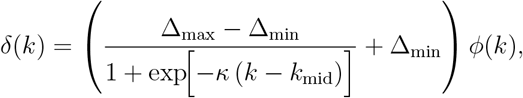

with parameters Δ_max_ = 0.75, Δ_min_ = 0.35, *κ* = 0.08, *k*_mid_ = 35, and

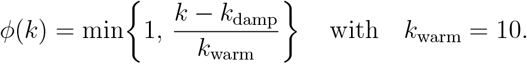

The state update is modified accordingly by

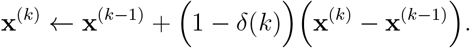

Second, a *network memory* mechanism is implemented to simulate the biological retention of past signals. The updated state is blended with the previous state via

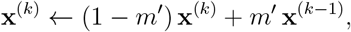

where *m*^*′*^ is an effective memory parameter (for example, *m*^*′*^ = 0.7 *m*). Finally, the state is non-linearly scaled to ensure biologically realistic values:

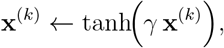

with *γ* = 1.2.

### 2.5 Convergence Criteria

The iterative process terminates when either the relative change in the state vector falls below a tolerance *ε* or when the maximum number of iterations *T* is reached. Specifically, convergence is determined by the condition

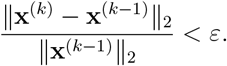

### 2.6 Loss Function and Parameter Optimization

Model parameters are calibrated by minimizing a composite loss function defined as

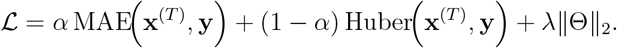

Here, **y** is the ground-truth perturbation vector, MAE represents the median absolute error, and the Huber loss is applied to manage outliers with a transition from quadratic to linear behavior beyond a threshold. The term Θ represents the set of model parameters (such as *σ, β, m, s, c*_1_, *c*_2_, *c*_3_, *α*_1_, *α*_2_, *α*_3_, etc.), and *λ* is the L2 regularization coefficient. Parameter optimization is performed using the Adam optimizer with early stopping based on validation error.

### 2.7 PETS Drug Efficacy Scoring

Upon convergence, the final state vector **x**^(*T*)^ is used to compute a drug efficacy (PETS) score, which quantifies the reversal of AD-related dysregulation. Let *E*_*i*_ denote the AD-specific expression change for protein *i* and *w*_*i*_ its corresponding importance weight. The PETS-score is computed as

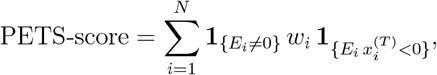

where **1**_{·}_ is the indicator function that equals 1 when the sign of 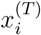 is opposite to the disease-induced change *E*_*i*_. To address potential off-target toxicity, pathway activation measures for toxicity-related gene sets can be further evaluated, using a penalty based on the severity of toxicity that is subtracted from the PETS-score to yield an adjusted efficacy metric.

### 2.8 Implementation and Visualization Pipeline

The full simulation pipeline is implemented in Python and proceeds through several sequential stages. First, the required data are loaded from CSV files, including the adjacency matrix **A**, drug vectors **d**, protein names, and auxiliary datasets. The adjacency matrix is normalized and the drug vector is standardized.

Next, multi-layer networks **A**^(1)^, **A**^(2)^, and **A**^(3)^ are computed and combined. Protein importance weights are integrated, and the simulation is then initialized with 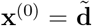 and iteratively updated, incorporating adaptive dampening, network memory, and non-linear scaling as described above.

After convergence, model parameters are fine-tuned by minimizing the composite loss function with the Adam optimizer and early stopping based on validation performance. Finally, the PETS-score is computed and time-series plots of protein activation trajectories are generated. The final activation states and scores are exported for downstream analysis.

### 2.9 Glossary of Key Terms

**Table 1:** Glossary of Key Terms Used in the Simulation Framework.

| Term | Definition |
| --- | --- |
| **Adaptive Dampening** | A mechanism that reduces the magnitude of state updates as the system approaches steady state to prevent oscillatory behavior. |
| **Network Memory** | The integration of historical state information into the current state to mimic biological memory effects. |
| **Boosting Vector** | An amplification factor applied to drug-affected nodes to enhance their contribution in the state update. |
| **Multi-Layer Propagation** | A method to incorporate higher-order interactions by raising the normalized adjacency matrix to successive powers and combining them. |
| **Convergence Criterion** | A predefined tolerance level that determines when the iterative simulation has sufficiently stabilized. |
| **Composite Loss Function** | A function that combines MAE, Huber loss, and regularization terms to calibrate model parameters. |

### 3 Results

### 3.1 PETS-Based Simulation of Alzheimer’s Disease Modulation

#### 3.1.1 Model Performance

The PETS algorithm was applied to an Alzheimer’s disease–relevant 501×501 protein–protein interaction (PPI) network. Initially, the model was trained using vorinostat—a well-characterized inhibitor of HDAC1, HDAC2, HDAC3, and HDAC6—via an iterative propagation scheme with gradient-based optimization. Convergence was achieved after more than 450 gradient descent iterations, yielding a final combined loss of 0.123, which corresponds to an average prediction error of approximately 6% relative to normalized SigCom L1000 data (Figure 2A).

**Figure 2.**
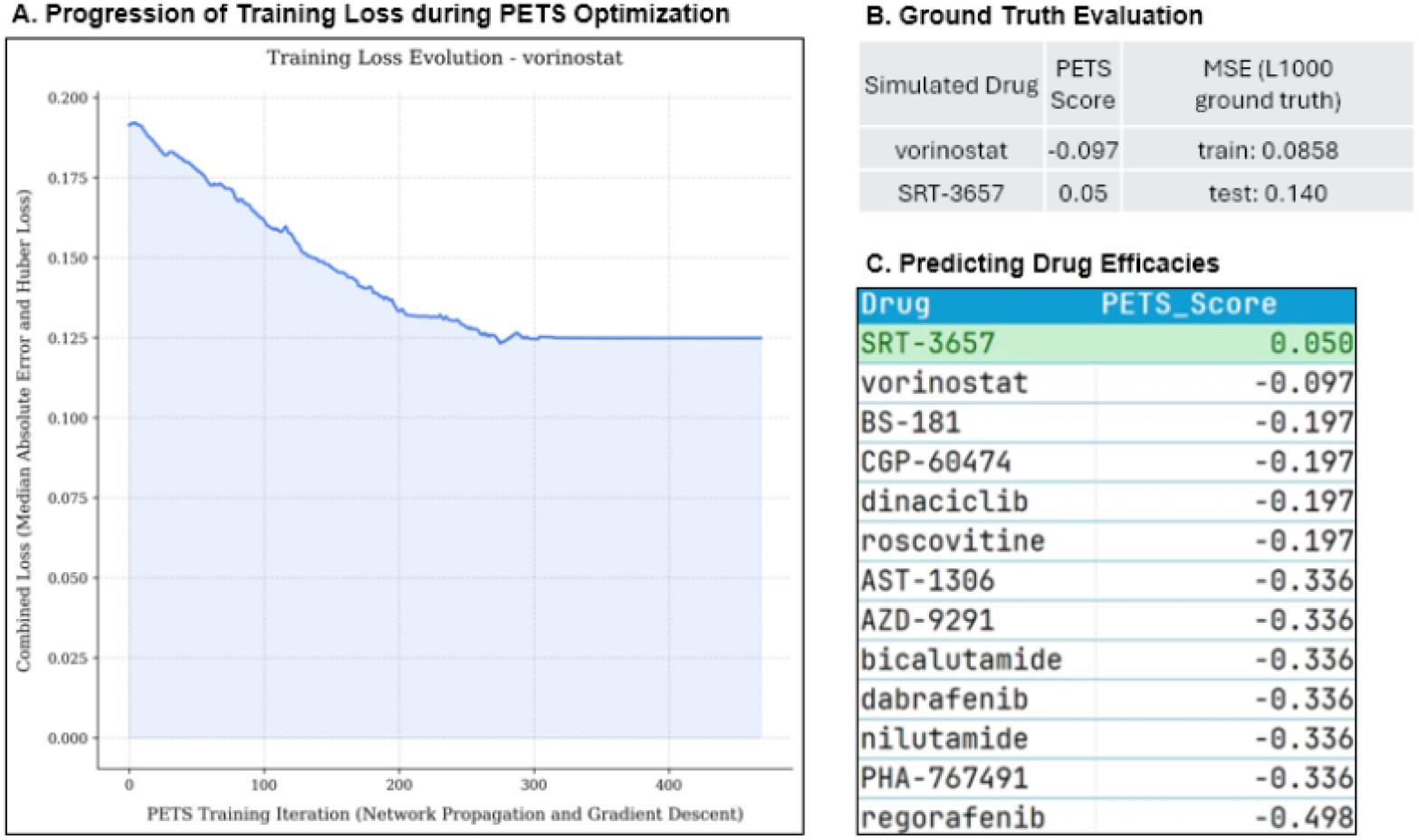
Performance and applications of the AD PETS model for simulating drug–gene perturbations. (A) The final training loss, computed as a weighted combination of median absolute error and Huber loss, converged to 0.123 after over 450 gradient descent iterations using vorinostat, demonstrating robust optimization. (B) PETS-simulated drug efficacy scores, along with their associated MSEs for both training and unseen test compounds, are benchmarked against SigCom L1000 ground truth data, underscoring the model’s predictive accuracy and generalizability. (C) A comparative evaluation of PETS scores across candidate compounds reveals that SRT-3657 achieves the highest efficacy score within our 501-gene iPS-ND34732 simulation framework, highlighting its therapeutic potential.

Subsequently, the model was evaluated using SRT-3657, a compound known to activate SIRT1. The PETS model predicted a drug efficacy score of 0.05 for SRT-3657. Validation against the SigCom L1000 dataset produced mean squared error (MSE) values of 0.0858 for training compounds and 0.140 for unseen test compounds, thereby demonstrating the model’s capacity to reliably simulate drug-induced perturbation profiles. Ground-truth gene expression data were preprocessed via quantile normalization followed by min–max scaling to ensure comparability across 12,327 genes.

#### 3.1.2 Experimental Consistency

We further evaluated the experimental consistency of the PETS simulations using iPS-ND34732 cells. For SRT-3657 treatment, simulated gene expression profiles for APP-related and tau-interacting genes were directly compared with corresponding experimental measurements. As shown in Figures 3a and 3b, regions with at least 75% concordance in both magnitude and sign relative to ground truth were classified as consistent; overall consistency improved with successive simulation iterations. Additionally, GeneTerrain visualizations—2D contour plot heatmaps of gene–gene expression networks—demonstrated close alignment between vorinostat-treated experimental cells and PETS-simulated profiles (Figures 3c and 3d), further validating the model’s capacity to recapitulate observed dynamics.

**Figure 3.**
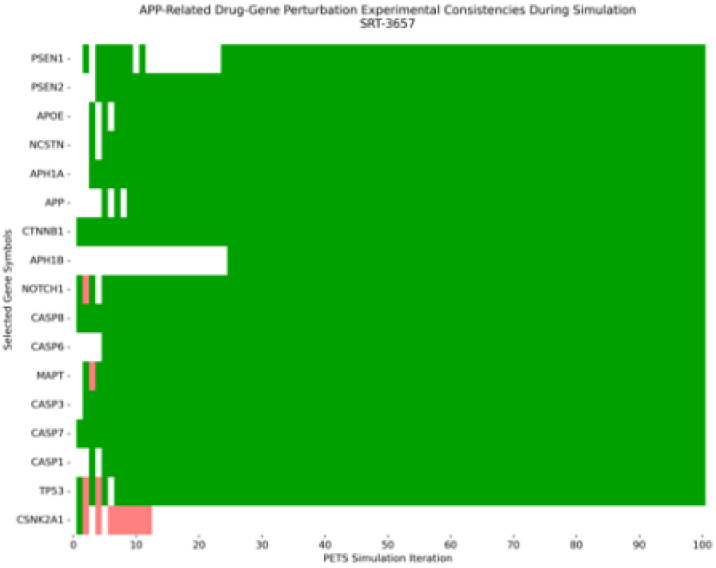
(a) Experimental consistency of simulated gene expression for APP-related genes under SRT-3657 treatment. Green bars indicate regions with ≥ 75% agreement in both magnitude and sign relative to ground truth; red bars denote inconsistencies; uncolored cells represent insufficient magnitude (*<* 75%).

**Figure 3b.**
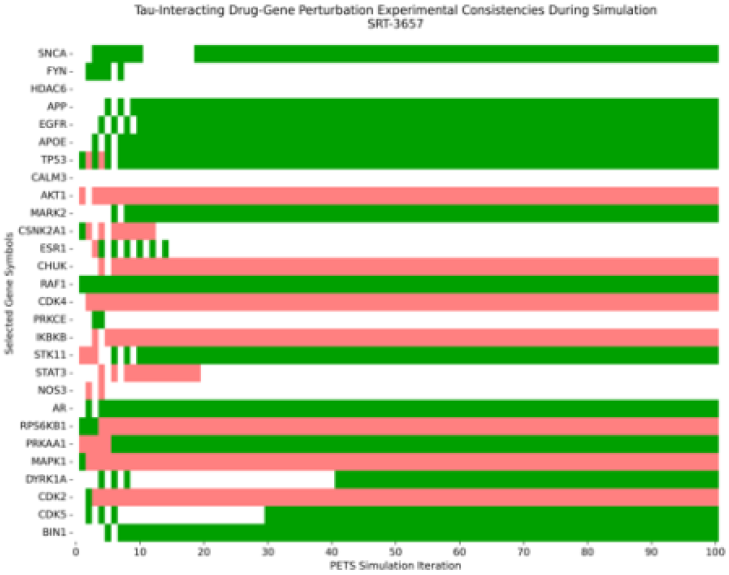
Alignment of simulated tau-interacting gene expression profiles with experimental data across PETS iterations. Gene sets were derived from PAG IDs TAX024082 and TAX026070, expanded using BEERE, and scored with the Rp metric via WINNER. Green and red regions denote consistent and inconsistent simulations, respectively.

**Figure 3c.**
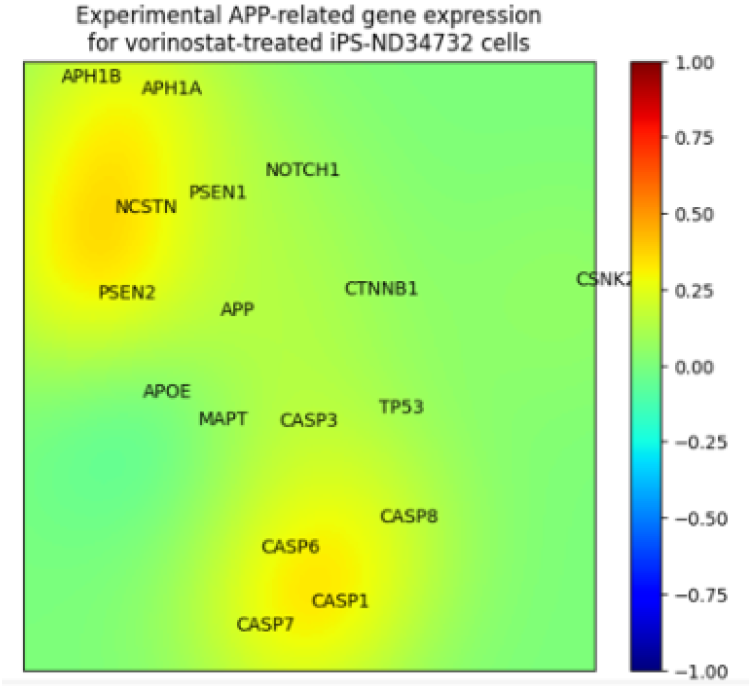
GeneTerrain visualization of APP-related effector gene expression in vorinostat-treated iPS-ND34732 cells. Expression levels are scaled to the range [−1, 1].

**Figure 3d.**
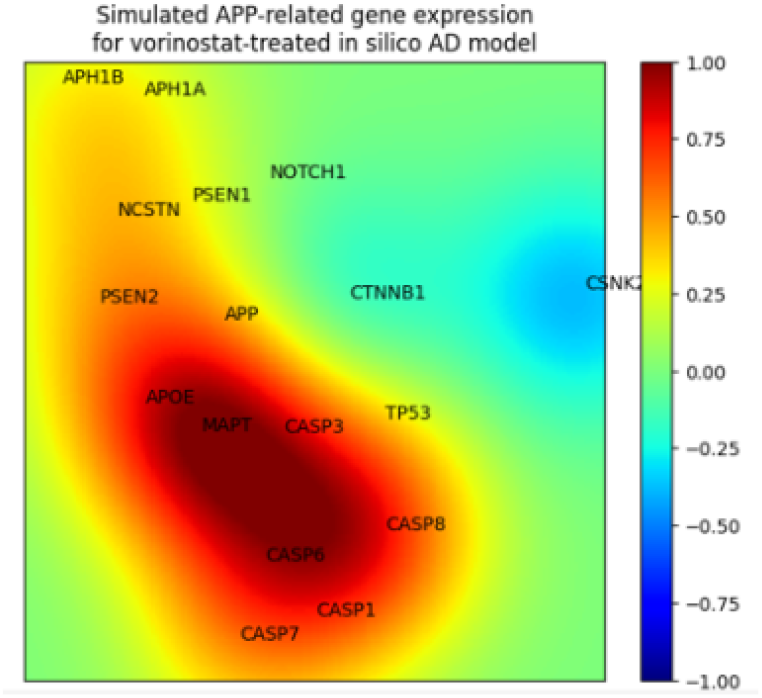
PETS-simulated GeneTerrain for APP-related effector genes. Moderate alignment with experimental terrain underscores the model’s ability to capture dynamic expression patterns.

### 3.2 Case Study: PETS Application to Alzheimer’s Disease Therapeutics Using a Large Interactome

To further illustrate the utility of the PETS framework in drug discovery for Alzheimer’s disease, we extended our analysis to a comprehensive interactome comprising 5,403 genes. Calibration was performed using high-fidelity ground truth from the SigCom L1000 dataset, which captures gene expression responses from SHSY5Y neuroblastoma cells treated with dasatinib. As shown in Figure 6, trajectories of the top upregulated and downregulated proteins converge and dampen over successive PETS iterations, reflecting robust model stability and effective parameter tuning.

Aggregate performance was quantified by comparing mean squared errors (MSEs) for 11 candidate compounds against the L1000 data (Figure 4). Concurrently, PETS-derived AD-gene reversal efficacy scores (Figure 5) provided an integrated measure of therapeutic potential. Our *in silico* analysis identified five compounds capable of modulating AD-related gene expression, with PHA-767491 emerging as the top candidate. This finding aligns with prior evidence suggesting that PHA-767491 may mitigate protein aggregation—a central pathological feature of Alzheimer’s disease [2].

**Figure 4.**
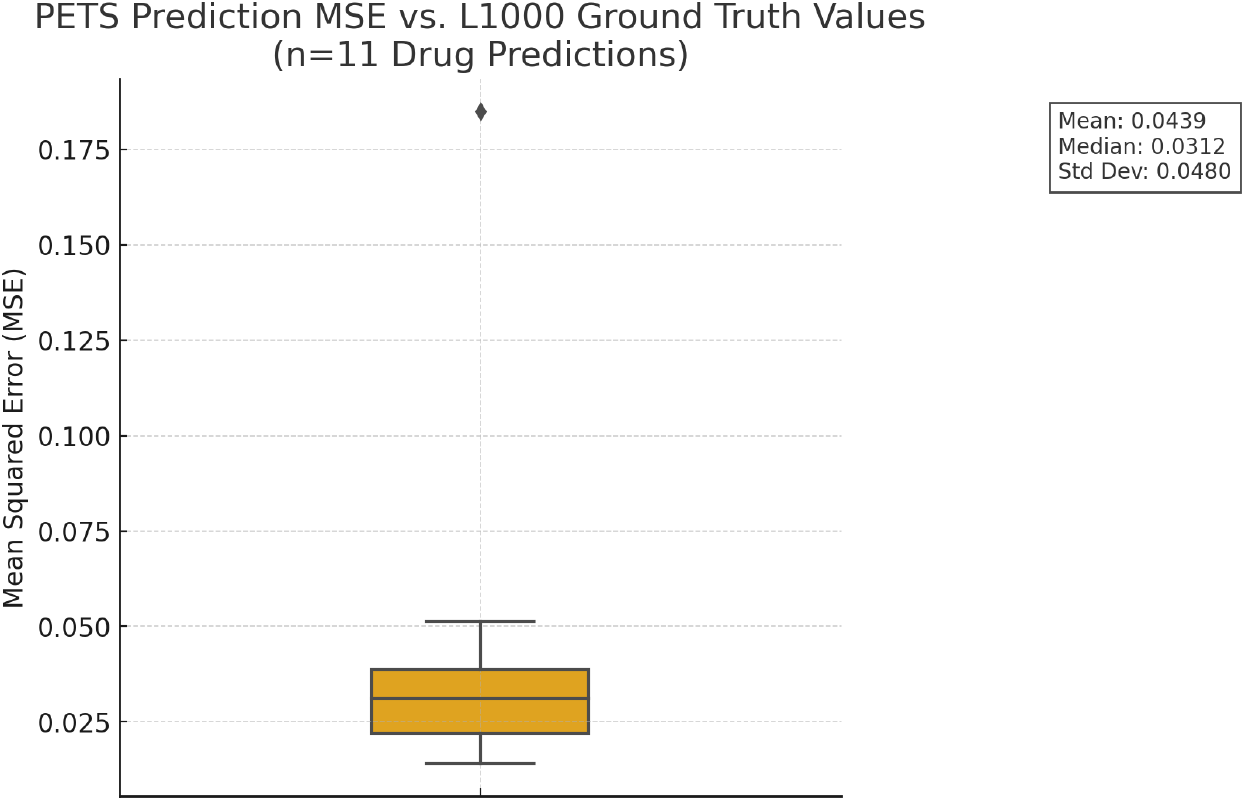
Aggregate mean squared errors (MSEs) for 11 candidate compounds benchmarked against the SigCom L1000 ground truth.

**Figure 5.**
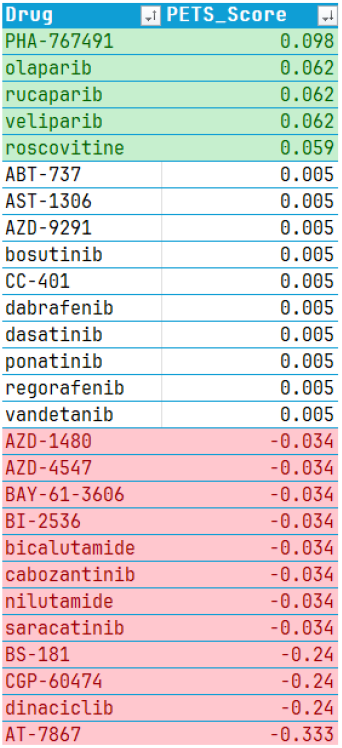
PETS-derived Alzheimer’s disease (AD) gene-reversal efficacy scores for 27 candidate compounds. PETS scores, derived from iterative simulations of gene regulatory network dynamics with robust normalization and differential expression analyses, quantify each compound’s ability to reverse AD-associated gene expression signatures. Notably, PHA-767491 achieved the highest reversal score, underscoring its potential as a promising AD intervention.

**Figure 6.**
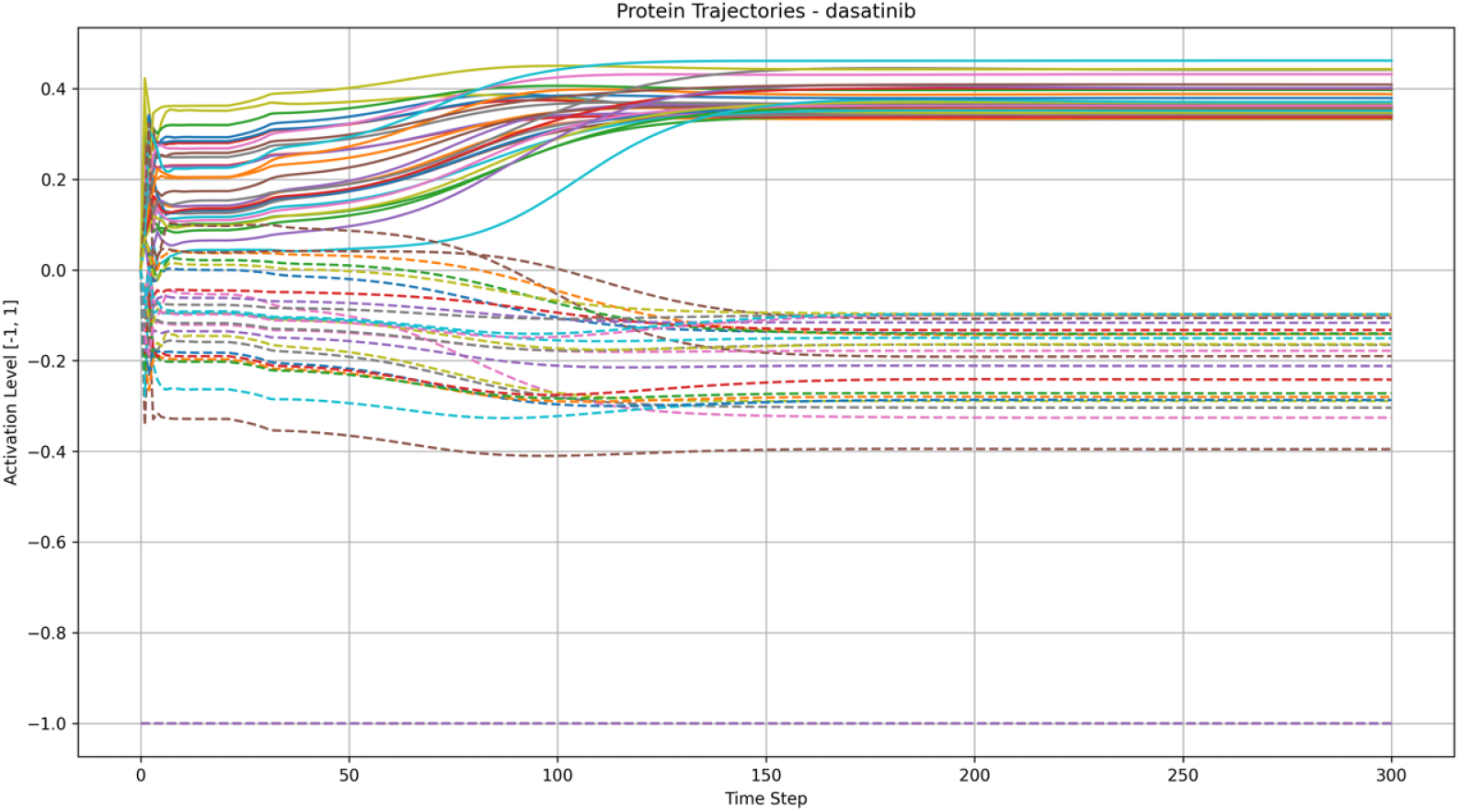
Convergence and dampening of the most upregulated and downregulated protein trajectories over successive PETS iterations in simulation of cells treated with dasatinib. Initial dynamic fluctuations in protein expression are progressively stabilized and attenuated over iterative PETS simulations. This behavior reveals the model’s capacity to capture inherent regulatory feedback loops and achieve a robust steadystate representation of dasatinib-induced proteomic perturbations, providing critical insights into the temporal evolution of protein network dynamics under therapeutic intervention.

Potential off-target toxicity was assessed by comparing gene expression profiles from our *in silico* AD model dosed with PHA-767491 to those from SHSY5Y cells treated with a hydrocinnamic acid negative control (PAG ID: TAX031886) from the L1000 dataset. Figure 7 illustrates a moderate upregulation of inflammatory and plaque-associated markers (including TLR4, TLR6, and CD36) following PHA-767491 treatment relative to control. Although these changes indicate some activation of toxicity pathways, their magnitude remains within acceptable limits, warranting further investigation into safety.

**Figure 7.**
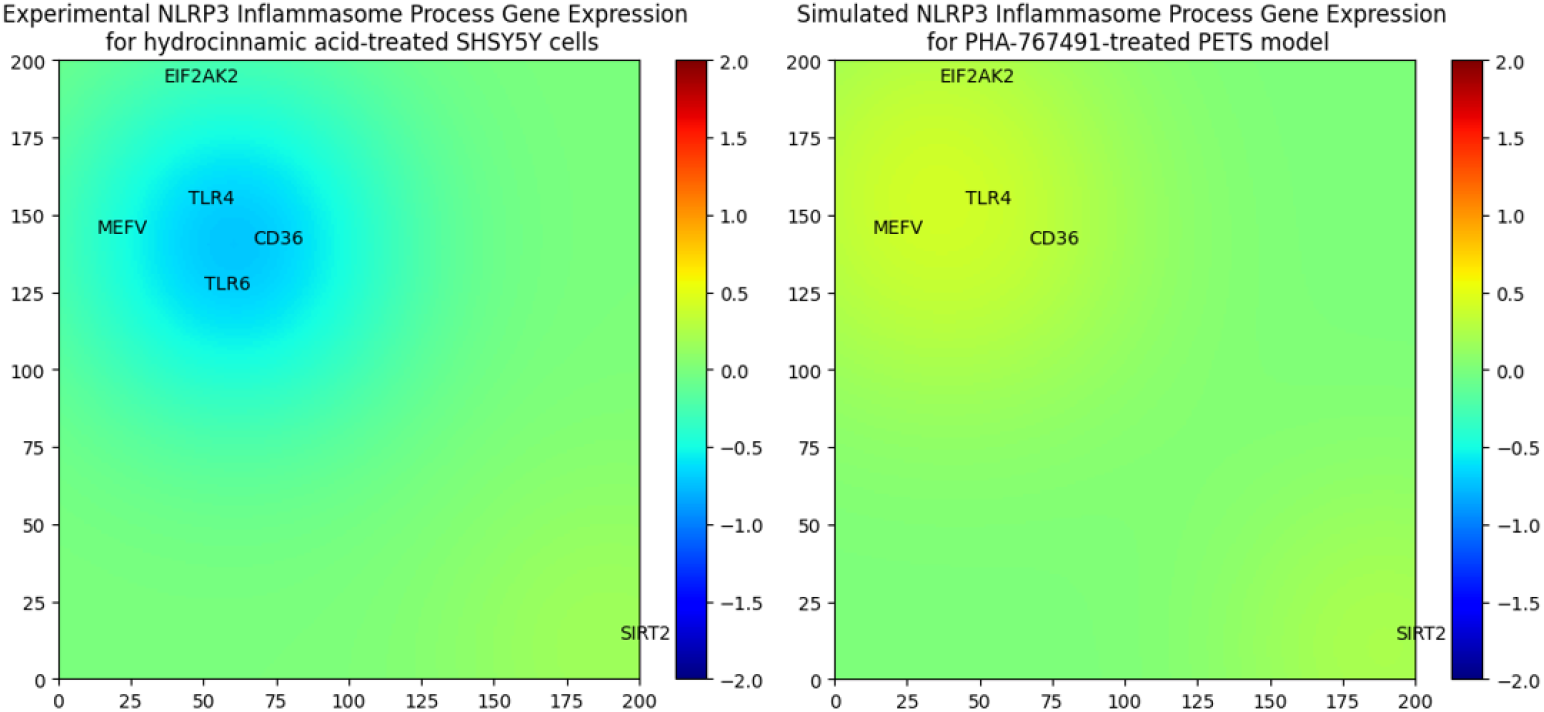
GeneTerrain comparison of expression profiles for NLRP3 inflammasome pathway genes. This figure juxtaposes the gene network expression landscapes of an *in silico* Alzheimer’s disease model dosed with PHA-767491 against SHSY5Y neuroblastoma cells treated with a hydrocinnamic acid negative control. The contour plot’s z-axis displays normalized z-scores computed from Level 3 SigCom L1000 log_2_ fold change data, processed via quantile normalization followed by min–max scaling. This rigorous pipeline preserves intrinsic transcriptional variability while ensuring robust comparability. The visualization reveals dynamic transcriptional perturbations elicited by PHA-767491, providing a quantitative benchmark against the negative control and yielding insights into the modulation of gene regulatory networks implicated in Alzheimer’s pathology.

The multi-scale simulation framework can also be employed to assess off-target toxicity by evaluating key pathways, including mitochondrial dysfunction (characterized by increased reactive oxygen species leading to cell death), cytochrome P450 interactions (which may result in liver damage), HERG potassium channel blockade (linked to sudden cardiac death), and COX-2 inhibition (associated with cardiovascular and gastrointestinal risks). By concurrently evaluating therapeutic efficacy (i.e., reversal of dysregulated AD expression) and off-target safety profiles, the PETS framework prioritizes compounds with robust disease-modifying potential and minimal safety risks.

### 3.3 Benchmarking Against Alternative Approaches

In addition to Alzheimer’s disease, the PETS algorithm was applied to a glioblastoma multi-forme (GBM) cell line study. Here, PETS-simulated gene expression profiles for dabrafenib-treated cells were compared to predictions from the GEARS algorithm [40]. Across nine dosage levels, PETS achieved lower median MSE values and narrower interquartile ranges compared to GEARS (Figure 8), underscoring its superior alignment with L1000 experimental data and more consistent predictive performance.

**Figure 8.**
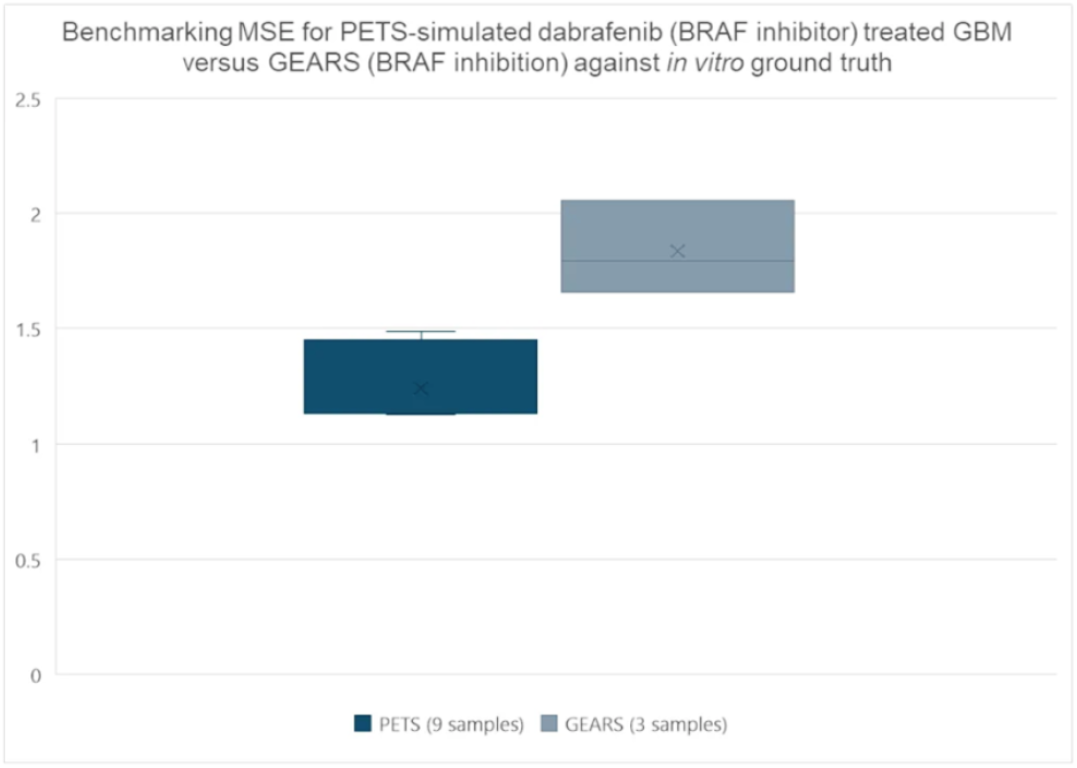
Benchmarking the PETS and GEARS frameworks in predicting gene expression responses in dabrafenib-treated glioblastoma (GBM) cells. Using L1000 experimental data across nine dosage levels, this analysis evaluates model performance based on key predictive metrics. PETS consistently achieved significantly lower median MSEs and narrower interquartile ranges (IQRs) compared to GEARS, highlighting its superior accuracy and precision in modeling dose-dependent transcriptional dynamics. These results demonstrate that PETS more reliably captures complex perturbations induced by BRAF inhibition, offering a robust computational tool for precision oncology through improved simulation of gene regulatory networks.

Our results demonstrate that the PETS algorithm effectively simulates drug-induced perturbations in both Alzheimer’s disease and GBM contexts. Its capacity to predict the reversal of AD-specific gene expression while simultaneously assessing off-target toxicity risks underscores its potential as a robust quantitative tool for precision medicine and drug repur-posing. Future work will focus on integrating dynamic network topologies, additional omics layers, and patient-specific data to further enhance predictive performance.

## 4 Discussion

The PETS framework represents a significant methodological advancement in simulating drug-induced perturbations, merging rigorous systems biology with state-of-the-art network modeling techniques. Our results demonstrate that by incorporating multi-layer representations of protein–protein interactions and adaptive dampening, PETS not only reproduces known pharmacodynamic responses but also reliably predicts the reversal of disease-specific drug–gene perturbation signatures.

A key strength of PETS is its ability to integrate both proximal and distal network effects via multi-layer propagation. By explicitly computing second- and third-order interactions, the framework captures cascading drug perturbation effects and elucidates complex feedback loops that conventional methods often overlook. Incorporation of protein importance weights (i.e., Rp scores) further refines predictions by emphasizing key regulatory nodes, which is critical for detecting subtle yet therapeutically relevant changes.

The iterative state update mechanism, augmented with adaptive dampening and memory incorporation, ensures numerical stability and convergence despite inherent oscillatory dynamics in multi-scale biological networks. In addition, the application of non-linear scaling aligns simulation outputs with experimentally observed calibrator dynamics, thereby enhancing predictive reliability.

Our case studies in Alzheimer’s disease and glioblastoma multiforme underscore the practical utility of PETS. In the Alzheimer’s model, the framework accurately reproduced gene expression reversals and identified candidate compounds such as PHA-767491 with promising therapeutic profiles, while concurrently assessing potential off-target toxicities. Benchmarking against established methods such as GEARS revealed that PETS achieves lower median mean squared errors (MSEs) and more consistent performance across multiple dosage levels. These findings position PETS as a robust platform for both hypothesis generation and candidate prioritization in drug repurposing pipelines.

Nonetheless, several limitations merit discussion. The current implementation is based on static, curated interactomes and expression profiles, which may not fully capture the dynamic nature of cellular responses in vivo. Furthermore, while the adaptive dampening strategy effectively suppresses oscillations, additional refinement is required to optimize convergence rates in highly heterogeneous networks. Addressing these challenges is essential for future iterations of the framework.

Overall, the PETS framework offers a blend of mechanistic insight and predictive power, serving as a critical tool for precision drug repurposing and bridging the gap between computational predictions and clinical outcomes.

## 5 Conclusions and Future Directions

We have introduced PETS—a novel multi-scale simulation framework that integrates curated pathway models with dynamic network propagation, adaptive dampening, and network memory—to predict drug–gene perturbations with high accuracy. Our results demonstrate that PETS can effectively reverse disease-specific dysregulation in complex interactomes, as evidenced in both Alzheimer’s disease and glioblastoma models, while also providing critical insights into potential off-target toxicity.

Looking forward, several avenues will further enhance the capabilities and impact of the PETS framework:

- **Integration of Multi-Omics and Patient-Specific Data:** Incorporating transcriptomic, proteomic, and metabolomic datasets will allow for the development of more comprehensive and personalized disease models, enhancing the predictive power of the framework.
- **Dynamic Network Topologies:** Adapting the model to incorporate time-varying and context-dependent network structures will better capture the evolving nature of cellular responses, particularly in chronic and multifactorial diseases.
- **Expansion to Combination Therapies:** Extending PETS to simulate multi-drug interactions and synergistic effects is a critical next step, enabling the exploration of combinatorial treatment strategies that address complex disease mechanisms.
- **Enhanced Validation and Clinical Translation:** Further validation against largescale clinical datasets, along with integration of electronic health record analytics and survival analysis, will be essential to translate PETS predictions into actionable clinical insights.
- **Algorithmic and Computational Optimizations:** Utilization of high-performance computing and advanced optimization techniques will facilitate the scaling of PETS to larger, more complex interactomes while reducing computational overhead.

These advancements will establish PETS as an indispensable tool in precision drug repurposing, guiding the discovery and clinical implementation of therapeutics with enhanced efficacy and minimized toxicity.

## 6 Code Availability

All underlying code, along with sample datasets and reproducible scripts, will be made openly available via GitHub (github.com/aimed-lab), ensuring that other researchers can adapt PETS to new disease contexts and replicate all analytical steps reported here.

## Acknowledgments

Special thanks to Thahn Nguyen and coauthors on the original PETS work. This work was supported by the University of Alabama at Birmingham and computational resources provided by the Systems Pharmacology AI Research Center.

